# Polysaccharide quantification using microbial enzyme cocktails

**DOI:** 10.1101/2024.07.10.602880

**Authors:** Sammy Pontrelli, Uwe Sauer

## Abstract

Polysaccharide quantification plays a vital role in understanding ecological and nutritional processes in microbes, plants, and animals. Traditional methods hydrolyze these large molecules into monomers, but these approaches are restricted to chemically hydrolysable polysaccharides. Enzymatic degradation is a promising alternative but typically requires the use of characterized recombinant enzymes or microbial isolates that secrete enzymes. In this study, we introduce a versatile method that employs undefined enzyme cocktails secreted by individual microbes or complex environmental microbial communities for the hydrolysis of polysaccharides. We focus on colloidal chitin and laminarin as representative polysaccharides of ecological relevance. Our results demonstrate that colloidal chitin can be effectively digested with an enzyme cocktail derived from a chitin-degrading *Psychromonas sp.* isolate. Utilizing a 3,5-dinitrosalicylic acid reducing sugar assay or liquid chromatography–mass spectrometry for mono- and oligomers detection, we successfully determined chitin concentrations as low as 62 mg/L and 15 mg/L, respectively. This allows for effective monitoring of microbial chitin degradation. To extend the applicability of our method, we also leveraged complex, undefined microbial communities as sources of enzyme cocktails capable of degrading laminarin. With this approach, we achieved a detection limit of 30 mg/L laminarin through the reducing sugar assay. Our findings highlight the potential of utilizing enzyme cocktails from both individual microbes and, notably, from undefined microbial communities for polysaccharide quantification. This advancement addresses limitations associated with traditional chemical hydrolysis methods.

## Introduction

Biological polysaccharides play vital roles in various functions, such as as energy storage molecules^1^, structuring cell walls^2,3^ and biofilms^4^, and supplying carbon to microbial communities^5–8^. Quantifying these polysaccharides is essential for studying their biological processes, yet their diverse chemical properties pose challenges in developing universally applicable quantification methods. The predominant methodologies for polysaccharide quantification involve hydrolyzing these complex molecules into monosaccharides, which are then quantified using colorimetric or chromatographic techniques^6,9–11^. Typically, this hydrolysis is conducted through chemical or enzymatic methods. One major limitation of chemical hydrolysis is its inefficacy across different polysaccharides, which may yield varying results or remain non-hydrolyzed^6^. Additionally, the harsh conditions employed in acid hydrolysis can damage acid-sensitive components^12,13^. A notable example of a polysaccharide resistant to acid hydrolysis is chitin, a linear homopolymer of β-(1,4)-linked N-acetylglucosamine (GlcNAc) monomers^14^. Chitin is the main constituent of mollusk and arthropod exoskeletons in marine environments^15^, making it one of the ocean’s most prevalent polysaccharides. While chitin can theoretically be deacetylated into chitosan to facilitate acid hydrolysis to glucosamine^16^, this approach is time-consuming and not suitable for microtiter plate formats.

On the other hand, enzymatic degradation presents a viable alternative for quantifying polysaccharides ^10,17–19^, especially those resistant to chemical hydrolysis. Enzymes produce specific degradation products that help identify particular polysaccharides in complex mixtures^17^. Nonetheless, a challenge in this approach lies in the difficulty of identifying enzymes with the desired activity. Moreover, enzyme expression and purification are time-consuming and may require optimization. An underutilized yet promising alternative is the utilization of microbial enzyme cocktails. This approach involves cultivating a polysaccharide-degrading bacterium or community on the target polysaccharide, which induces the expression of enzymes with desired hydrolytic activities. The resulting undefined mixture of secreted enzymes, referred to as a cocktail, can be concentrated from the supernatant and subsequently used to digest a polysaccharide followed by quantifying mono- or oligomers. Previous studies have demonstrated the application of enzyme cocktails to determine monomeric compositions of polysaccharides^20,21^, as well as to evaluate the catalytic capabilities^21–23^ and relative activities ^21,22^ of secreted enzymes. For instance, purified enzyme cocktails derived from a characterized fucoidan-degrading microbe have successfully been used for the degradation and quantification of fucoidan^20^, indicating the potential of enzyme cocktails as a method for polysaccharide quantification.

We present a methodology for developing enzyme cocktail-based assays to quantify polysaccharides, extending this approach to utilize natural microbial communities. This effectively addresses the limitations of chemical hydrolysis and the reliance on recombinant enzymes or specific degrading microbes. As proof of concept for quantifying colloidal chitin, we extracted an enzyme cocktail from the model chitin-degrading isolate *Psychromonas sp. Psy*6C06^24,25^. Building on this foundation, we used undefined microbial seawater communities to induce a laminarin-degrading enzyme cocktail, which we then purified to quantify laminarin. This method offers a readily adaptable solution for quantifying polysaccharides across various biological systems.

## Methods

### Chemicals and strains

All chemicals were obtained from Sigma Aldritch unless otherwise specified.

### Preparation of colloidal chitin

To prepare colloidal chitin as described^22,26^, 10 grams of powdered chitin from shrimp shells were dissolved in 100 mL of concentrated phosphoric acid and incubated at 4°C for 48 hours. After incubation, 500 mL of milliQ water were added to precipitate the chitin, which was then purified by vacuum filtration. The chitin was dialyzed against water for 72 hours using a cellulose dialysis tubing membrane (Sigma D9402) to remove any remaining phosphoric acid and subsequently resuspended in water to achieve a concentration of approximately 15 g/L. The suspension was homogenized using a Bosch SilentMixx Pro blender to obtain a uniform colloidal chitin solution. To determine the exact chitin concentration, a 10 mL aliquot was vacuum-concentrated overnight, and the dried chitin was weighed. Based on this measurement, the original suspension was diluted to obtain a final concentration of 10 g/L chitin. The colloidal chitin was autoclaved before use as a bacterial substrate to ensure sterility.

### LC-MS quantification

Measurements were performed using Liquid Chromatography (Agilent Infinity II UHPLC stack) coupled with an Agilent 6520 Time of Fight Quadrupole Time of Flight Mass Spectrometer in negative mode, 4GHz, high resolution mode.

For quantification of chitin oligomers, an Agilent EC-CN Poroshell column (50×2.1 mm, 2.7 µM) was used isocratically to reduce interference of salts on metabolite ionization^27^. The buffer contained 90% water, 10% (v/v) acetonitrile (CHROMOSOLVE) and 0.1% (v/v) formic acid. The flow rate was 0.35 mL/min. Prior to measurement, the sample was diluted 20-fold in millQ water. 3 µL was injected every 2 minutes. Data analysis and quantification was performed using Agilent Quantitative Masshunter software.

For time course laminarin pentose degradation, an Agilent EC-CN Poroshell column (50×2.1 mm, 2.7 µM) was used isocratically as described above. Here, the assay was initiated in the Agilent Infinity autosampler by adding 50 µL of 10 mM laminarin pentose to 50 µL of enzyme cocktail, and 3 µL of the enzyme reaction was injected every 3 hours. Data analysis was performed through manual peak integration using Agilent Qualitative Masshunter software.

For quantification of laminarin oligomers of different chain lengths, an Agilent HILIC-Z Poroshell column (100×2.1 mm, 1.8 µM) was used. Buffer A contains 10% (v/v) acetonitrile and 0.3% (v/v) ammonium hydroxide. Buffer B contains 90% acetonitrile (v/v) and 0.3% ammonium hydroxide (v/v). The flow was 0.9 mL/min and the column was kept at 35°C. The gradient is as follows. 0% A for 1 minute, followed by a gradient to 40% A until 5 minutes. This is held for 1 minute before dropping to 0% A and re-equilibration for 3 minutes. 3 µL of sample was injected. Data analysis and quantification was performed using manual peak integration with Agilent Qualitative Masshunter software.

### Enzyme cocktail purification

*Psy*6C06 was streaked from −80°C glycerol stocks onto 1.5% (w/v) agar plates of MB 2216 (Fisher) medium. A single colony was used for an overnight preculture in MB 2216, and the following day the preculture was inoculated 1% into 400 mL of simplified MBL medium with 2 g/l colloidal chitin and shaken at 200 rpm until early stationary phase. Simplified MBL medium is created from several stocks: 4-fold concentrated seawater salts (NaCl, 80 g/L; MgCl_2_*6H_2_O, 12 g/L; CaCl_2_*2H_2_O, 0.6 g/L; KCl, 2 g/L), 1000-fold diluted sodium sulfate (1 M), 500-fold diluted phosphate dibasic (0.5mM), and 20-fold diluted HEPES buffer (1 M pH 8.2).

After growth, the medium was centrifuged at 2800 relative centrifugal force for 20 minutes to remove any remaining chitin and cells, then sterile filtered using a 0.2 µM membrane. This supernatant was then concentrated 10-fold using an Amicon stirred cell (Millipore) with a 3 kDa cutoff filter. A protease inhibitor was added to the final concentrate (Roche cOmplete EDTA free protease inhibitor cocktail). 500 µL aliquots were added to 1.5 mL microcentrifuge tubes, which were then snap frozen by placing them in liquid nitrogen before storing at −80°C until use. The concentration of protein in the chitinase enzyme cocktail is 0.07 mg/mL as quantified using Bradford reagent.

For generation of laminarinases from seawater microbes, the same growth and purification protocol was followed, except the medium contained 1 g/l Laminarin from *Laminaria digitata* (Sigma) as the carbon source and 10 mM ammonium chloride as the nitrogen source. Seawater was collected from the North Sea on the shore of Cuxhaven (53°53’04.3“N 8°37’57.1”E), passed through a 5 µM filter, mixed 1:1 with 80% (v/v) glycerol, and stored at −80 °C until inoculation. The concentration of protein in the laminarinase enzyme cocktail is 0.012 mg/mL as quantified using Bradford reagent.

### Chitin quantification assay

Frozen enzyme aliquots were thawed on ice. 100 µL of enzyme was added to 100 µL of sample and placed at room temperature until sampling. If the sample is suspected to contain more than 500 mg/L chitin, it was diluted to fall within this quantifiable range. To sample, the assay was centrifuged at 2800 rcf for 1 minute and the supernatant was collected and stored at −20°C until enzyme products are measured using DNS reagent or LC-MS.

### Laminarin quantification assay

Frozen enzyme aliquots were thawed on ice. For laminarin oligomer formation, 50 µL of enzyme was added to 50 µL of laminarin pentamer (Megazyme, dissolved in 40 mM ammonium bicarbonate buffer pH 7.8) and placed at room temperature for 24 hours. The reaction mixture was directly quantified using LC-MS as described above. For laminarin quantification, 50 µL laminarin from *Laminaria digitata* was added at various concentrations to 50 µL of enzyme cocktail. Samples were taken at 24 hours and directly quantified using DNS reagent.

### DNS reagent preparation

100 mL of water was heated to 70-75°C, then poured over 2 g of NaOH pellets with constant stirring. 2.18 g of 3,5-dinitrosalicylic acid was then dissolved in this mixture, and 30 g of sodium potassium tartrate (Rochelle salts) were added. The solution was cooled to room temperature and stored in the dark.

### DNS reducing sugar assay

25 µL of sample was added to 75 µL of DNS reagent and the mixture was heated to 95°C for 15 minutes in a thermocycler. 80 µL was added to a 384 well plate and a colorimetric readout was measured at OD_540_.

## Results

### Assay design

To cultivate a polysaccharide-degrading microbe or complex microbial community, the organism(s) is grown on the target polysaccharide until it reaches early stationary phase. This process selects for degraders and induces the expression of hydrolytic enzymes. Following cultivation, the cells are filtered to obtain a cell-free supernatant that contains the secreted enzymes. These enzymes are then concentrated tenfold using a 3kDa molecular weight cutoff filter, aliquoted, snap-frozen in liquid nitrogen, and stored at −80°C. When ready for use, the enzyme cocktail is thawed and mixed with a polysaccharide sample to initiate hydrolysis. After hydrolysis, the supernatant is collected for the quantification of monomers and oligomers using either a reducing sugar assay or liquid chromatography-mass spectrometry (LC-MS). Akin to other enzyme-based quantification methods^17,28^, this assay relies on consistent enzyme activity across all samples, leading to the production of uniform sets of oligomers and monomers. This consistency establishes a quantitative relationship between the amount of polysaccharide digested and the number of reducing sugar ends and degradation products formed. A calibration curve is calculated based on digestion of standards with known polysaccharide concentrations. These standards are digested concurrently with unknown samples to account for variations in enzyme activity among different cocktail batches, any degradation that may occur during storage, or residual mono- or oligomers that may remain in the cocktails after concentration.

### Quantification of colloidal chitin using enzyme cocktails from a bacterial isolate

We evaluated the assay design for quantifying colloidal chitin using an enzyme cocktail obtained from the chitin-degrading *Psychromonas sp.* bacterium *Psy*6C06, which encodes 13 chitinases^22^. To confirm chitinase activity, the cocktail was incubated for 72 h with a sample containing 2 g/L of colloidal chitin at equal volumes. The resulting mono- and oligomers were analyzed by LC-MS, detecting both monomeric GlcNAc and oligomers of various chain lengths. Notably, the primary degradation products observed were monomeric GlcNAc and dimeric chitobiose, consistent with previous characterizations of the chitinases secreted by *Psy*6C06^22^ **(Fig. 2A)**. To assess the time required for the cocktail to completely hydrolyze samples containing up to 2 g/L of colloidal chitin, we measured the time-resolvedformation of reducing sugar ends using the 3,5-dinitrosalicylic acid (DNS) reagent **(Fig. 2B)**. Within 72 hours, complete hydrolysis was achieved only in samples with 500 mg/L colloidal chitin or less, as evidenced by a stable signal. Consequently, all subsequent experiments were standardized to a 72-hour hydrolysis period, with samples known or suspected to contain more than 500 mg/L colloidal chitin diluted prior to digestion.

**Figure 1:**
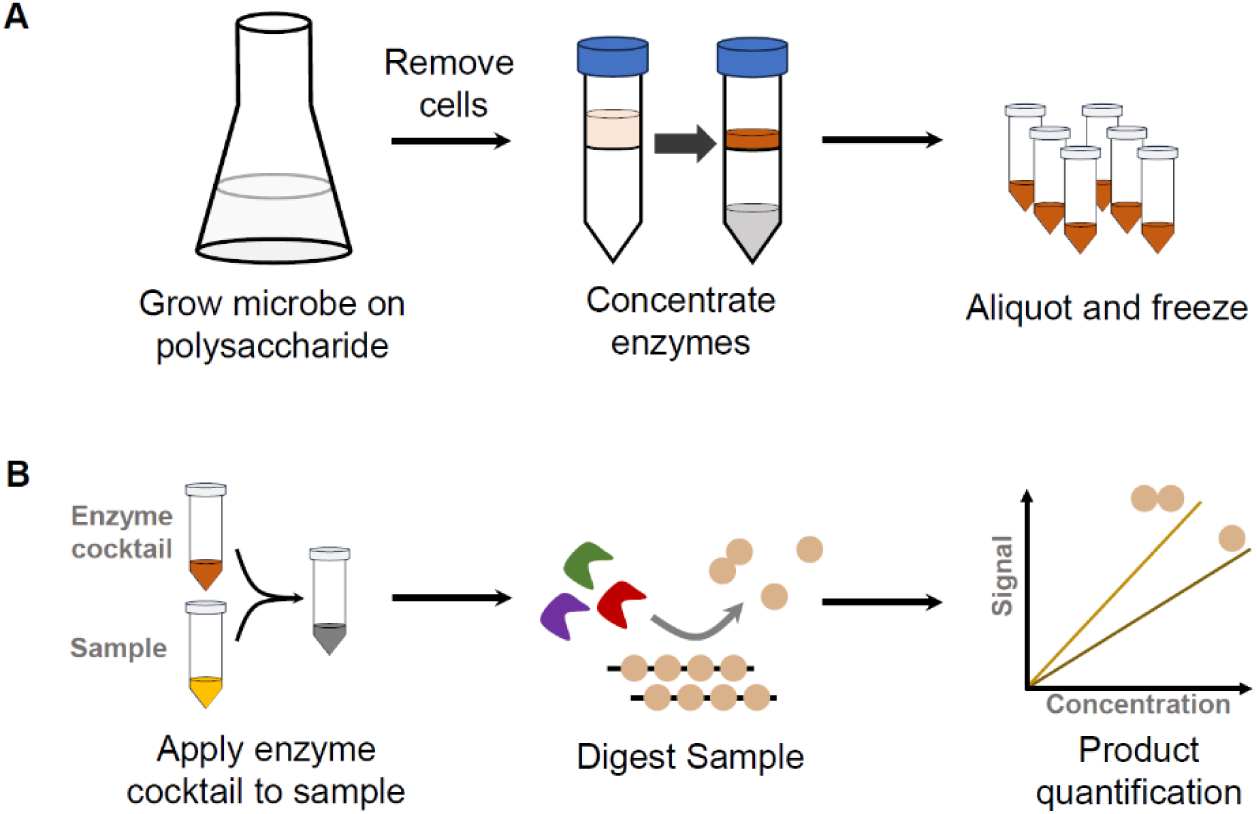
Assay design and workflow. A) Enzyme cocktail preparation involves growing a microbe, or microbial community, on the polysaccharide as a sole carbon source, followed by filtering cells and concentrating enzymes using a 3 kDa molecular weight cutoff membrane. The concentrate is aliquoted and stored until use. B) Polysaccharide quantification entails mixing the enzyme cocktail with a sample to initiate digestion, leading to the formation of monomers or oligomers. Post-digestion, the products can be quantified using LC-MS, HPLC, chemical, or other methods.

**Figure 2:**
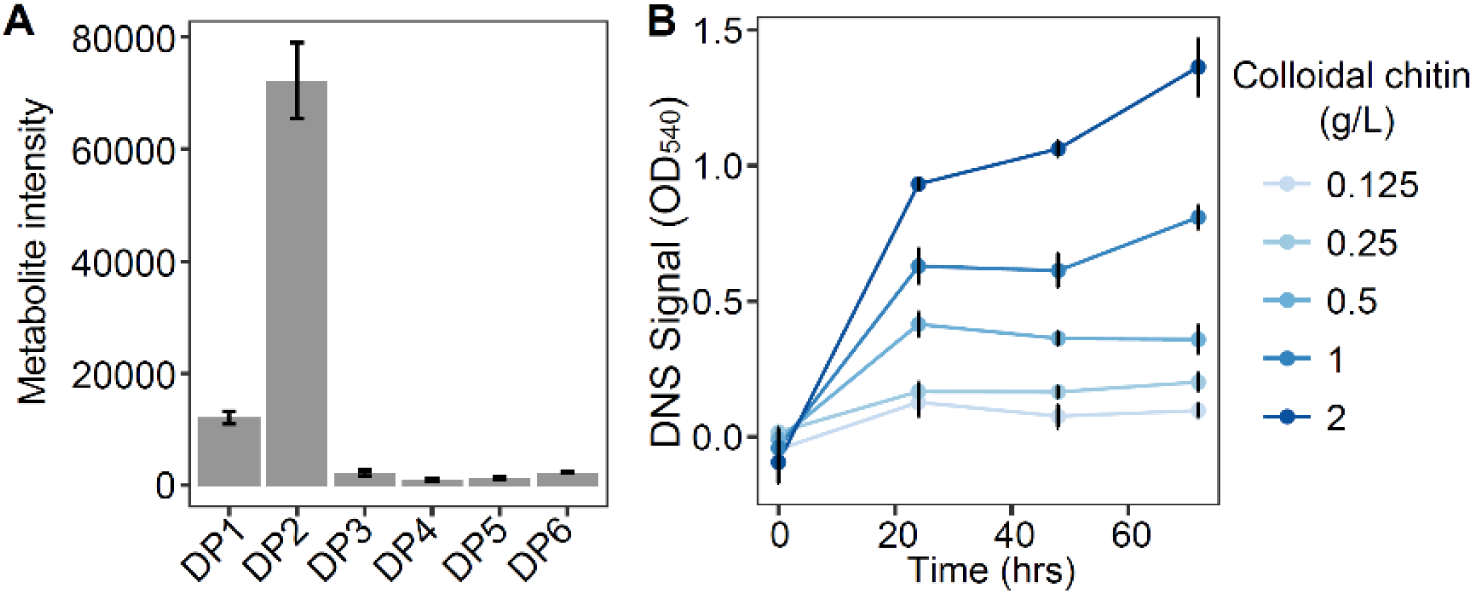
Activity of chitin enzyme cocktail. A) GlcNAc oligomers (Degree of polymerization, DP 1-6) formed after 72 hr digestion of 2 g/L colloidal chitin by the enzyme cocktail, measured using LC-MS. B) Samples with varying concentrations of colloidal chitin digested with equal volumes of enzyme cocktail. Liberation of mono- and oligomers is measured every 24 hours for 3 days using a DNS reducing sugar assay (with an OD_540_ signal). Experiments were performed in triplicate. Error bars represent standard deviation from the mean.

Enzyme cocktails often comprise multiple hydrolytic enzymes that can degrade not only the target polysaccharide, but also other polysaccharides present in a sample. Consequently, if an alternative polysaccharide is non-specifically degraded into mono- and oligomers, colorimetric assays like the DNS assay, which reacts with various reducing sugar ends, may lead to an overestimation concentrations of the target polysaccharide. In contrast, techniques such as LC-MS or HPLC can effectively distinguish between different mono- and oligomers after sample digestion, making these degradation products valuable for specific readouts of the target polysaccharide degradation. As previously demonstrated^17,28^, this distinction is particularly beneficial when the enzyme digestion generates known, specific oligomers from the target polysaccharide, allowing for the identification of a specific polysaccharide’s oligomers even amidst a mixture of multiple polysaccharides. Here, the enzyme cocktails from *Psy*6C06 primarily yield GlcNAc and chitobiose as the major degradation products, both of which serve as effective readouts for quantifying chitin. However, in cases where it is uncertain whether other GlcNAc-containing polysaccharides are present in a sample that might undergo non-specifically hydrolyzed, quantifying chitin based on chitobiose provides a viable alternative. This is due to chitobiose being confirmed as a specific product of chitin enzyme digests because direct GlcNAc-GlcNAc linkages are generally uncommon in natural polysaccharides. We compared the use of chitobiose measured by LC-MS as a readout against two other readouts: GlcNAc measured with LC-MS and total reducing sugar ends measured with the DNS reagent. We defined the limit of detection (LOD) as the lowest concentration yielding a signal above the background mean plus five standard deviations. We obtained an LOD of 15 mg/L colloidal chitin when using chitobiose as a readout, compared to 500 mg/L with GlcNAc **(Fig. 3ABC)**, and 62.5 mg/L when using DNS. These results demonstrate that using chitobiose as a readout not only enhances the specificity of chitin quantification but also provides the most sensitive detection among the evaluated methods.

**Figure 3:**
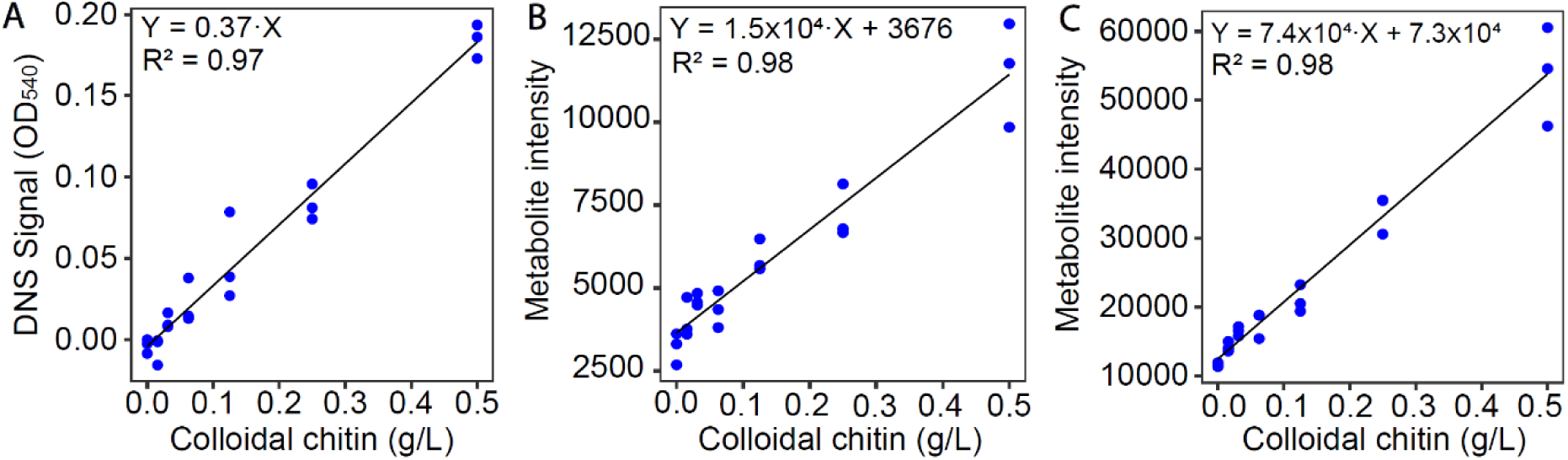
Assay sensitivity. Decreasing concentrations of colloidal chitin are applied to enzyme cocktail and digested for 72 hours. A) Detection of total reducing sugar ends with the DNS reagent. B) Detection of GlcNAc by LC-MS. C) Detection of dimeric chitobiose by LC-MS. A linear regression of signal intensity per chitin concentration is shown.

### Measuring colloidal chitin dynamics in bacterial cultures

Quantifying the dynamics of chitin degradation is important for investigating bacterial cultures and the role of marine communities in biogeochemical carbon cycling, as it enables to correlate changes in population dynamics with chitin degradation rates. Using the above enzyme cocktail, we monitored the time-course consumption of colloidal chitin by the chitin-degrading *Vibrio splendidus Vib*1A01^29,30^. During growth of *Vib*1A01, we periodically sampled aliquots, centrifuged them, and subjected the pellet to brief boiling to denature any chitinases derived from the organism. This denatured pellet was then incubated with the enzyme cocktail at a 1:1 ratio for 72 h. To ensure that other components in the *Vib*1A01 biomass were not non-specifically hydrolyzed by the enzyme digest, we digested biomass from a *Vib*1A01 culture grown on 20 mM GlcNAc instead of chitin. This test confirmed the absence of any detectable background signal using the DNS reagent (data not shown). Our results show a gradual decrease in colloidal chitin that corresponded with the growth of *Vib*1A01 **(Fig. 4)**. This finding highlights the method’s effectiveness in quantifying chitin degradation in bacterial cultures, which has broader implications for researchers investigating chitin dynamics in various biological systems.

**Figure 4:**
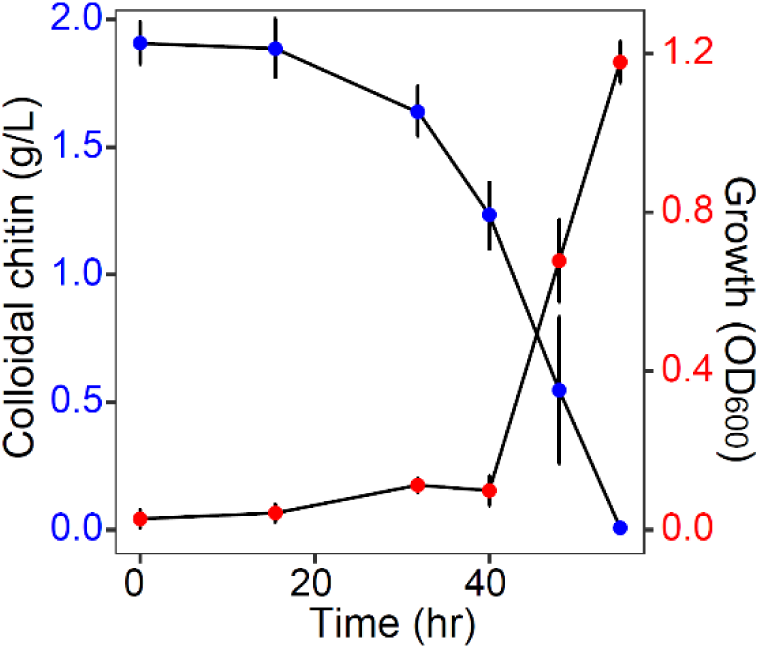
Chitin concentrations during bacterial growth. OD_600_ of chitin degrader *Vib*1A01 when growing on chitin as a sole carbon source (Red). Colloidal chitin concentration at each of the measured timepoints (Blue) measured using DNS reagent Error bars represent standard deviation of the mean of three biological replicates.

### Laminarin quantification using enzyme cocktails from environmental communities

When degrading microbes are unavailable or cannot be isolated from the environment, an alternative is to collect enzyme cocktails from complex environmental microbial communities. By growing these communities on the polysaccharide of interest as a sole carbon source, specific degraders will be selected, and their secreted enzymes can be concentrated into a cocktail. We tested this approach for quantifying laminarin, an algal storage polysaccharide with a β-(1-3)-glucose backbone and single β-(1-6)-glucose side branches. Seawater collected from the North Sea near the shore of Cuxhaven (53°53’04.3“N 8°37’57.1”E) was used to inoculate a medium containing laminarin as the sole carbon source, thereby inducing and concentrating laminarinases withing the generated enzyme cocktail. To assess the activity of this cocktail, we digested 5 mM of a β-(1-3)-glucose pentamer, which represents the laminarin backbone, for four hours. We measured the formation of smaller oligomers using LC-MS, which revealed a mixture of monomers and smaller oligomers **(Fig. 5A)**, indicating the presence of multiple laminarinases with varying activities. Further, we determined the time required for the complete digestion of 2 mM laminarin pentamer (equivalent to 1.6 g/L) using LC-MS, observing a stable signal before 21 hours **(Fig. 5B)**. Consequently, we used 24 hours as an appropriate digestion time for laminarin concentrations of up to 1 g/L. To corroborate that the DNS reagent could also be employed to quantify laminarin, offering a more widely accessible measurement method beyond LC-MS, we used it to detect laminarin concentrations between 1 g/L and 0.03 g/L after a 24-h digestion period, achieving a LOD of 30 mg/L **(Fig. 5C)**. These results underscore the potential of environmental microbial communities as a valuable source of enzyme cocktails for quantifying various polysaccharides.

**Figure 5:**
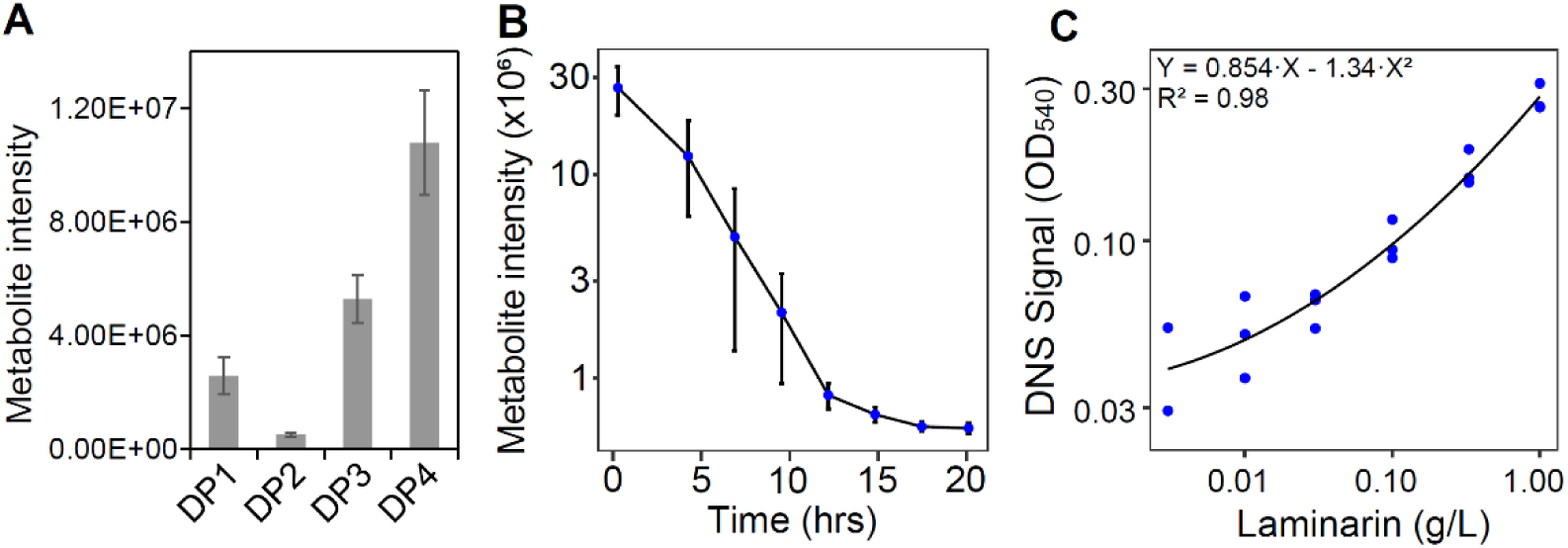
Laminarin quantification with seawater community enzymes. A) Mass spectrometry measurement of glucose oligomers (Degree of polymerization, DP1-4) after 4 hour digestion of a laminarin pentamer by a laminarinase enzyme cocktail expressed by a seawater microbial community. B) Intensity of laminarin pentamer measured with LC-MS during time-course digestion with an enzyme cocktail, starting from an initial 2mM concentration. C) Digestion of different laminarin concentrations by an enzyme cocktail, quantified with DNS reagent and a readout of OD_540_. A polynomial regression is shown and the LOD is 30 mg/L. Error bars represent standard deviation of the mean of three biological replicates.

## Discussion

In this study, we demonstrate that microbial enzyme cocktails can be sourced from both complex microbial communities and individual isolates to quantify polysaccharides, specifically using colloidal chitin and laminarin as case studies. Unlike other enzyme-based techniques for polysaccharide quantification, our approach eliminates the need for identifying and recombinantly expressing specific hydrolytic enzymes. Instead, we collect these enzymes from cultures of bacteria that are actively degrading the target polysaccharide. By concentrating all secreted enzymes into single cocktails, we also retain potential non-catalytic proteins that may enhance the activities of hydrolytic enzymes^31^. Furthermore, we show that enzyme cocktails can be derived from complex microbial communities, thus circumventing the need to isolate specific polysaccharide degrading microbes.

A limitation of this approach is the possibility of cocktails containing other hydrolytic enzymes that may degrade alternative polysaccharides in complex samples, which could inflate the quantified values. One strategy to address this issue is to conduct control experiments to eliminate background interference. For example, we showed that the *Psy*6C06 enzyme cocktail does not degrade polysaccharides in *Vib*1A01 biomass, thereby ensuring no interference when measuring the degradation of colloidal chitin. Furthermore, LC-MS can be utilized to monitor specific mono- and oligomers, rendering the quantification of target polysaccharides more reliable, particularly when the cocktail produces polysaccharide-specific oligomers, such as chitobiose. However, LC-MS techniques that rely solely on exact mass identification may encounter challenges when non-specifically degraded polysaccharides share similar monosaccharide compositions. This complication is particularly relevant for laminarin, wich is entirely composed of glucose, making its quantification difficult in the presence of other glucans and glucanases, such as glycogen with its α(1→4) glucose backbone and α(1→6) branches. To overcome this challenge for improved accuracy, employing LC-MS using MS^2^ capabilities can be advantageous. This technique resolves monosaccharide linkages in the released oligomers that are polysaccharide specific.

This work introduces an innovative approach for developing quantification assays using enzyme cocktails derived from complex microbial communities. By leveraging these communities, we eliminate the need to identify and purify recombinant enzymes or depend on characterized microbial isolates. Our method requires only a microbial isolate or community that secretes hydrolytic enzymes, making it broadly applicable to a variety of polysaccharides that are degraded by extracellular enzymes, and facilitating the efficient use of natural microbial communities as sources of enzyme. Importantly, we applied this approach to quantify colloidal chitin—a particulate polysaccharide that is resistant to chemical hydrolysis—thereby addressing a significant methodological gap in the study of this abundant marine polysaccharide^22,32–34^. Our approach offers a valuable tool for investigating microbial degradation of specific polysaccharides in controlled environments. It provides insights into the physiology and behavior of microbes, as well as precise analysis of their roles in biopolymer turnover. This research ultimately advances the understanding of polysaccharide utilization in microbial ecology, biotechnology, and biogeochemical cycles.

## Acknowledgements

We greatly acknowledge financial support from the Simons Foundation through the Principles of Microbial Ecosystems (PriME) collaboration (grant 542395).

## Bibliography

1. Suzuki, E. & Suzuki, R. Variation of Storage Polysaccharides in Phototrophic Microorganisms. J. Appl. Glycosci. 60, 21–27 (2013).

2. Lerouxel, O., Cavalier, D. M., Liepman, A. H. & Keegstra, K. Biosynthesis of plant cell wall polysaccharides — a complex process. Current Opinion in Plant Biology 9, 621–630 (2006).

3. Hemsworth, G. R., Déjean, G., Davies, G. J. & Brumer, H. Learning from microbial strategies for polysaccharide degradation. Biochemical Society Transactions 44, 94–108 (2016).

4. Raszka, A., Chorvatova, M. & Wanner, J. The role and significance of extracellular polymers in activated sludge. Part I: Literature review. Acta hydrochim. hydrobiol. 34, 411–424 (2006).

5. Cerqueira, F. M., Photenhauer, A. L., Pollet, R. M., Brown, H. A. & Koropatkin, N. M. Starch Digestion by Gut Bacteria: Crowdsourcing for Carbs. Trends in Microbiology 28, 95–108 (2020).

6. Arnosti, C. et al. The Biogeochemistry of Marine Polysaccharides: Sources, Inventories, and Bacterial Drivers of the Carbohydrate Cycle. Annu. Rev. Mar. Sci. 13, 81–108 (2021).

7. Enke, T. N. et al. Modular assembly of polysaccharide-degrading marine microbial communities. Current Biology 29, 1528–1535.e6 (2019).

8. Lindemann, S. R. A piece of the pie: engineering microbiomes by exploiting division of labor in complex polysaccharide consumption. Current Opinion in Chemical Engineering 30, 96–102 (2020).

9. Decamp, A. et al. A New, Quick, and Simple Protocol to Evaluate Microalgae Polysaccharide Composition. Marine Drugs 19, 101 (2021).

10. Schiavone, M. et al. A combined chemical and enzymatic method to determine quantitatively the polysaccharide components in the cell wall of yeasts. FEMS Yeast Res 14, 933–947 (2014).

11. Rühmann, B., Schmid, J. & Sieber, V. High throughput exopolysaccharide screening platform: From strain cultivation to monosaccharide composition and carbohydrate fingerprinting in one day. Carbohydrate Polymers 122, 212–220 (2015).

12. Lin, P. & Guo, L. Spatial and vertical variability of dissolved carbohydrate species in the northern Gulf of Mexico following the Deepwater Horizon oil spill, 2010–2011. Marine Chemistry 174, 13–25 (2015).

13. Cao, X. et al. A Major Step in Opening the Black Box of High-Molecular-Weight Dissolved Organic Nitrogen by Isotopic Labeling of *Synechococcus* and Multibond Two-Dimensional NMR. Anal. Chem. 89, 11990–11998 (2017).

14. Peter, M. G. Applications and Environmental Aspects of Chitin and Chitosan. Journal of Macromolecular Science, Part A 32, 629–640 (1995).

15. Keyhani, N. O. & Roseman, S. Physiological aspects of chitin catabolism in marine bacteria11This paper is publication 521 from the McCollum-Pratt Institute. Biochimica et Biophysica Acta (BBA) - General Subjects 1473, 108–122 (1999).

16. Yan, X. & Evenocheck, H. M. Chitosan analysis using acid hydrolysis and HPLC/UV. Carbohydrate Polymers 87, 1774–1778 (2012).

17. Becker, S., Scheffel, A., Polz, M. F. & Hehemann, J.-H. Accurate Quantification of Laminarin in Marine Organic Matter with Enzymes from Marine Microbes. Appl Environ Microbiol 83, e03389–16 (2017).

18. Steinke, N., Vidal-Melgosa, S., Schultz-Johansen, M. & Hehemann, J. Biocatalytic quantification of α-glucan in marine particulate organic matter. MicrobiologyOpen 11, e1289 (2022).

19. Sun, Y., Li, L., Zhang, Y., Xue, C. & Chang, Y. An enzyme-pHBH method for specific quantification of porphyran. International Journal of Biological Macromolecules 257, 128530 (2024).

20. Buck-Wiese, H. et al. Fucoid brown algae inject fucoidan carbon into the ocean. Proc. Natl. Acad. Sci. U.S.A. 120, e2210561119 (2023).

21. Sichert, A. et al. Verrucomicrobia use hundreds of enzymes to digest the algal polysaccharide fucoidan. Nat Microbiol 5, 1026–1039 (2020).

22. Pontrelli, S. et al. Metabolic cross-feeding structures the assembly of polysaccharide degrading communities. Sci. Adv. 8, eabk3076 (2022).

23. Unfried, F. et al. Adaptive mechanisms that provide competitive advantages to marine bacteroidetes during microalgal blooms. The ISME Journal 12, 2894–2906 (2018).

24. Ebrahimi, A., Schwartzman, J. & Cordero, O. X. Cooperation and spatial self-organization determine rate and efficiency of particulate organic matter degradation in marine bacteria. Proc. Natl. Acad. Sci. U.S.A. 116, 23309–23316 (2019).

25. Enke, T. N., Leventhal, G. E., Metzger, M., Saavedra, J. T. & Cordero, O. X. Microscale ecology regulates particulate organic matter turnover in model marine microbial communities. Nature Communications 9, 2743 (2018).

26. Pontrelli, S. et al. Competition strategies driving resource partitioning in chitin degrading communities. Preprint at 10.1101/2024.11.07.622309 (2024).

27. Pontrelli, S. & Sauer, U. Salt-tolerant metabolomics for exometabolomic measurements of marine bacterial isolates. Anal. Chem. 93, 7164–7171 (2021).

28. Deng, Y., Chen, L.-X., Zhu, B.-J., Zhao, J. & Li, S.-P. A quantitative method for polysaccharides based on endo-enzymatic released specific oligosaccharides: A case of Lentinus edodes. International Journal of Biological Macromolecules 205, 15–22 (2022).

29. Iffland-Stettner, A. et al. A Genome-Scale Metabolic Model of Marine Heterotroph Vibrio splendidus Strain 1A01. mSystems 8, e00377–22 (2023).

30. Amarnath, K. et al. Stress-induced metabolic exchanges between complementary bacterial types underly a dynamic mechanism of inter-species stress resistance. Nature Communications 14, 3165 (2023).

31. Vaaje-Kolstad, G., Horn, S. J., van Aalten, D. M. F., Synstad, B. & Eijsink, V. G. H. The Non-catalytic Chitin-binding Protein CBP21 from Serratia marcescens Is Essential for Chitin Degradation. Journal of Biological Chemistry 280, 28492–28497 (2005).

32. Szabo, R. E. et al. Historical contingencies and phage induction diversify bacterioplankton communities at the microscale. Proc. Natl. Acad. Sci. U.S.A. 119, e2117748119 (2022).

33. Shen, C.-R., Chen, Y.-S., Yang, C.-J., Chen, J.-K. & Liu, C.-L. Colloid Chitin Azure Is a Dispersible, Low-Cost Substrate for Chitinase Measurements in a Sensitive, Fast, Reproducible Assay. SLAS Discovery 15, 213–217 (2010).

34. Salas-Ovilla, R., Gálvez-López, D., Vázquez-Ovando, A., Salvador-Figueroa, M. & Rosas-Quijano, R. Isolation and identification of marine strains of *Stenotrophomona maltophilia* with high chitinolytic activity. PeerJ 7, e6102 (2019).

